# High frequency electrical stimulation entrains fast spiking interneurons and bidirectionally modulates information processing

**DOI:** 10.1101/2025.07.01.662561

**Authors:** Pierre Fabris, Eric Lowet, Krishnakanth Kondabolu, Yangyang Wang, Yuxin Zhou, Xue Han

## Abstract

**Background:** Clinical intracranial electrical stimulation often deploys trains of high frequency pulses. While brief bursts of stimulation are known to heterogeneously modulate neuronal spiking, it is unclear how trains of high frequency pulses influence neural dynamics.

**Objective:** As fast spiking interneurons (FSIs) can support rapid firing, we seek to determine how high frequency stimulation modulates FSIs.

**Methods:** We characterized the real-time effect of one-second-long local stimulation at 40 versus 140 Hz on parvalbumin positive interneurons, known as FSIs, in motor and visual cortices in awake mice using near kilohertz voltage imaging, free of electrical stimulation artifact.

**Results:** Stimulation at 140 Hz, like 40 Hz, heterogeneously modulates individual FSIs membrane voltage in both cortices, leading to complex temporal dynamics. FSIs in both cortices are robustly entrained by 40 Hz stimulation, even though 40 Hz led to prominent membrane hyperpolarization in visual cortex but not motor cortex. Intriguingly, visual cortical FSIs, but not motor cortical ones, were reliably entrained by 140 Hz stimulation. Finally, while stimulation consistently reduced the response amplitude of visual cortical FSIs to visual flickers, response temporal precision is bidirectionally modulated.

**Conclusion:** High frequency electrical stimulation mediates brain-region specific entrainment of FSIs, and bidirectionally modulates FSI temporal processing of synaptic inputs. Thus, high frequency stimulation can differentially engage inhibitory neurons in different brain regions to modulate network information processing.

**Highlights:**

- Evoked membrane potential (Vm) responses are frequency and brain region specific
- 140 Hz stimulation entrains the Vm of visual, but not motor, cortical FSIs
- 140 Hz, but not 40 Hz, is effective at reducing Vm amplitude to visual flickers
- Stimulation bidirectionally modulates Vm response timing to visual inputs
- Visual cortical FSIs are suppressed by 40 Hz stimulation, unlike other conditions

## Introduction

Intracranial electrical stimulation is routinely used to treat various neurological disorders, including Parkinson’s Disease, essential tremor, dystonia, and epilepsy^1–5^. Across the broad parameter space of stimulation pulse patterns, frequency has been widely explored to tune therapeutic efficacy and to reduce side effects^2, 6–9^. In general, frequencies above 100 Hz tend to be more consistently effective than lower frequencies^2, 10^. However, the mechanisms underlying the distinct effects of stimulation frequencies remain unclear^6^. Single-pulse electrical stimulation led to diverse effects on neuronal firing in humans and preclinical models^11–15^, activating or suppressing^16^ neurons either directly at the site of stimulation, or indirectly through antidromic activation of axons to connected areas^17–21^. However, clinical neuromodulation generally delivers trains of electrical stimulation, often at high frequencies. Given the large stimulation artifacts on electrodes, the real time effect of stimulation pulse trains at different frequencies is much less understood.

Electrical stimulation serves as an external input to a neuron and thus influences neuronal processing of intrinsic synaptic inputs. It has been postulated that high frequency stimulation creates a functional informational lesion within targeted circuits^22–25^. A previous experimental attempt on testing this hypothesis using cellular voltage imaging demonstrated that electrical stimulation suppressed hippocampal neuronal responses to artificial optogenetic inputs^23^. In addition to response amplitude, another important consideration is response timing or temporal precision^26^, exemplified by the various oscillation features that have been linked to behavior and pathology^27–30^. As electrical stimulation could increase membrane conductance, passively reducing the amplitude of the evoked responses, neuromodulation may also bias cellular responses to more salient or larger synaptic inputs, leading to improved temporal precision to inputs.

Individual neuron’s response to electrical stimulation is heavily influenced by its intrinsic biophysical properties^31, 32^ and synaptic inputs^33^. Due to the limited membrane voltage (Vm) temporal response kinetics, most mammalian neurons cannot reliably support high frequency firing beyond a hundred hertz and thus serve as a low-pass filter for rapid electric field changes. For example, stimulation evoked firing rate is limited to ∼50-80 Hz for cortical pyramidal neurons^34,35^. However, some neurons, such as cerebella granule cells^36, 37^, cortical parvalbumin positive fast spiking interneurons (FSIs)^38, 39^, striatal medium spiny neurons^40^ can burst at frequencies well beyond 100 Hz. These neurons, capable of rapid spiking, may be better driven by high-frequency stimulation (>100 Hz). Indeed, high frequency stimulation has been shown to increase spiking in thalamic brain slices^41^. Similarly, cellular calcium dynamics in motor cortex FSIs versus non-FSIs were differentially modulated by high frequency intracranial electrical stimulation^42^ or transcranial ultrasound stimulation^43^. However, it remains unclear how Vm of FSIs, capturing both synaptic inputs and spiking output, respond to high frequency stimulation.

To characterize how high frequency stimulation modulates Vm of FSIs, we performed cellular voltage imaging from cortical neurons expressing FSI marker protein parvalbumin in awake mice.

We compared the Vm responses in FSIs in motor versus visual cortices, during one-second-long 140 versus 40 Hz local intracranial electrical stimulation. To further understand the effect of stimulation on cellular information processing, we characterized the impact of electrical stimulation on the response of visual cortex FSIs to visual flickers presented to awake mice.

## Results

### Intracranial electrical stimulation evokes diverse membrane voltage (Vm) changes across individual FSIs in both motor and visual cortices

To examine how electrical stimulation influences individual cortical FSIs, we performed high-speed cellular voltage imaging using the genetically encoded voltage sensor SomArchon^44^ in head-fixed mice freely locomoting. Briefly, under general anesthesia, we infused AAV9-Flex-SomArchon-GFP^44^ to transduce neurons expressing the FSI marker gene parvalbumin (PV) in the superficial layers of visual or motor cortices of PV-Cre mice. We then placed a glass window for chronic optical access to the transduced neurons, a pair of stainless electrodes across the glass window for local intracranial stimulation, and a metal bar for head fixation (Fig. 1A, B). Voltage imaging was performed at least 3 weeks post viral transduction to allow for sufficient SomArchon expression (Fig. 1Ci-Fi, and Fig. S1). During each recording session, SomArchon fluorescence from individual FSIs was recorded at about ∼830 frames/sec using custom microscopes while mice were awake, head fixed, voluntarily navigating a spherical treadmill (Methods).

**Figure 1:**
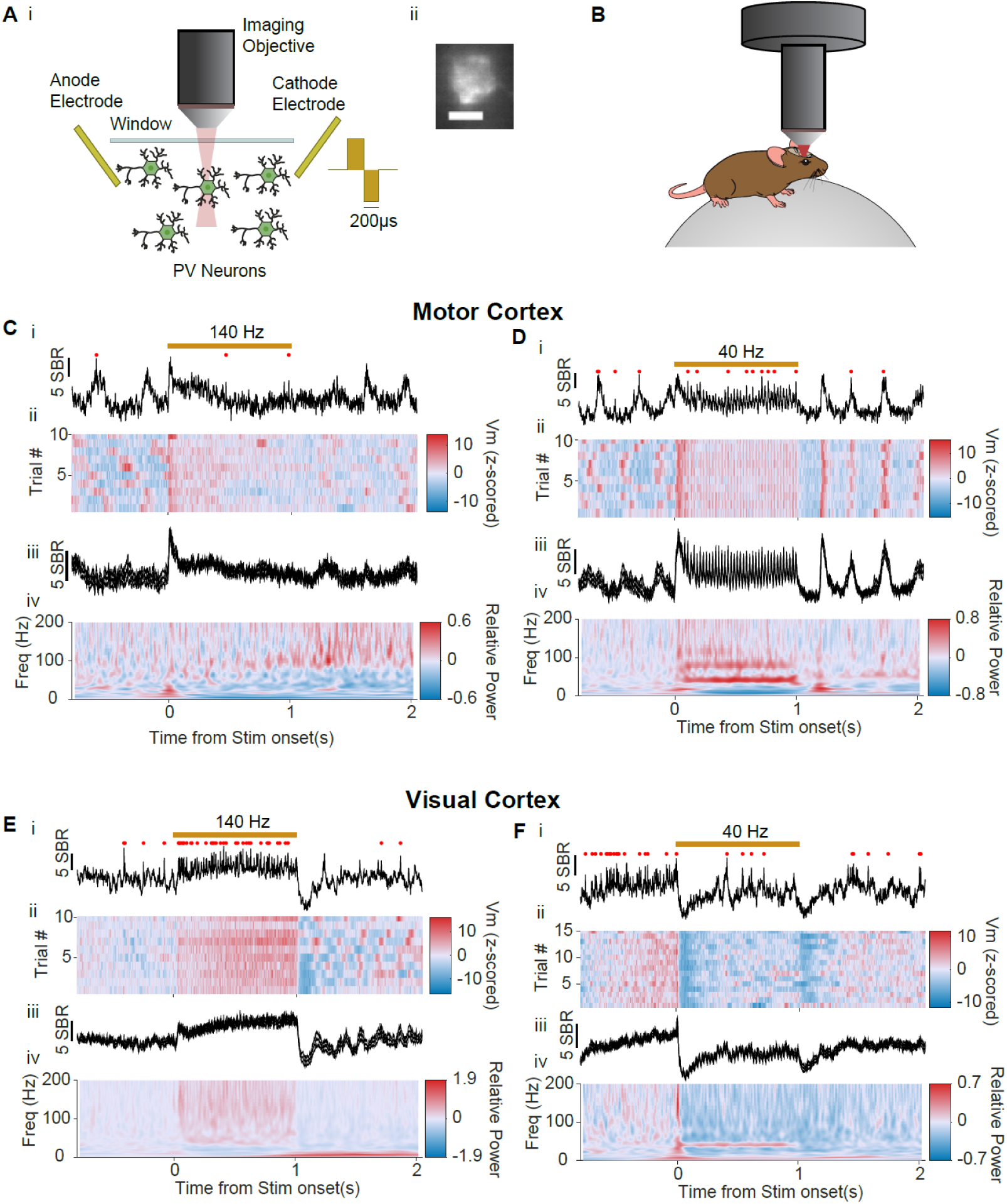
High speed cellular voltage imaging reveals the real-time effects of local intracranial stimulation on the membrane voltage (Vm) of cortical FSIs in awake mice. **(A)** (i) Illustration of optical voltage imaging of FSIs during local electrical stimulation. (ii) An example neuron expressing SomArchon (scale bar is 10μm). (B) Illustration of animal preparation and microscope setup. (C) Example Vm response of a motor cortex FSI during 140 Hz stimulation. (i) Single trial Vm trace with red markers indicating detected spikes. (ii) Heatmap of Vm across all trials aligned to stimulation onset. (iii) Trial-averaged Vm. Shaded area shows the SEM. (iv) Power spectra of Vm. (D) Same as c, but for an example motor cortex FSI during 40 Hz stimulation. (E) Same as c, but for a visual cortex FSI during 140 Hz stimulation. (F) Same as e, but for a visual cortex FSI during 40 Hz stimulation. Blue scale bars for Vm are 5 SBR (signal to baseline ratio).

Electrical stimulation was delivered across the imaging field of view at either 40 Hz or 140 Hz. Stimulation pulses were biphasic, 200 μs per phase, square-wave, and charge balanced (Fig. 1A). Stimulation current was tuned for each recording to account for variations across animal preparations and recording sites. Specifically, we first empirically determined the minimum current required to evoke a neural response at 40 Hz in each animal by gradually increasing current amplitude at 10 μA increments until a visible change in SomArchon fluorescence was detected. During subsequent recordings, the current amplitude was further adjusted to account for small variations in the relative distance between the imaging field of view and the implanted electrodes, using similar procedures starting from the minimum current of the given animal (Methods). All current used produced no noticeable behavioral changes.

Once an effective current was determined, we recorded 10-15 trials per neuron, with each trial containing 1 second baseline, 1 second stimulation, and 1 second post-stimulation, and an inter-trial-interval of 10 seconds. The recorded SomArchon image frames were motion corrected, visually inspected, and then manually segmented to identify individual neurons. SomArchon fluorescence for each neuron was then extracted and processed to obtain the cellular Vm traces and spikes (Fig. 1Ci-Fi, Methods).

Across motor cortex recordings, the stimulation current was similar for 140 Hz and 40 Hz (122.5±147.6 μA for 140 Hz, mean ± standard deviation, n=20 recordings from 5 mice, and 101.9±60.4 μA for 40 Hz, n=21 recordings from 4 mice) (Fig. S2A, B). Across visual cortex recordings, the stimulation current was also similar for 140 Hz (147.6±54.9 μA, n=42 recordings from 6 mice) and 40 Hz (167.4±58.7 μA, n=38 recordings from 6 mice) (Fig. S2A, C). However, visual cortex recordings generally had higher amplitude than motor cortex recordings for both 40 and 140 Hz stimulation (Fig. S2A), suggesting a brain region difference in sensitivity to stimulation. The current amplitude used stayed largely consistent across recording days (Fig. S2B, C).

We noted large variations in stimulation-evoked responses across neurons, with some exhibiting transient depolarization within 100 ms of stimulation onset (Fig. 1C) and others showing sustained depolarization throughout the entire stimulation period (Fig. 1D, E). While not as prevalent, some neurons showed prominent Vm hyperpolarization, which was sometimes proceeded by a brief depolarization (Fig. 1F). As previously reported in the mouse CA1^23^, most neurons were reliably entrained by 40 Hz stimulation, with Vm changes following stimulation pulses and exhibiting a prominent increase in 40 Hz Vm spectral power (Fig. 1Div, Fiv). Even though the Vm responses were heterogeneous across neurons (Fig. S3), stimulation evoked changes were largely consistent across trials within the same neuron (Fig. 1 Cii-Fii, Ciii-Fiii).

### While motor cortical FSIs are predominantly activated by stimulation at both frequencies, visual cortical FSIs are robustly suppressed by 40 Hz and activated by 140 Hz

To characterize the evoked responses in individual FSIs, we examined Vm amplitude change during 100 ms (transient period) or 100-1000 ms (sustained period) after stimulation onset, or Vm entrainment throughout the stimulation period. Briefly, to determine whether a neuron’s Vm amplitude was modulated, we computed the mean Vm during the transient and the sustained period for each trial and compared them to the mean Vm during the baseline period, defined as the 500 ms immediately before stimulation onset (Methods). Using Wilcoxon’s sign-rank test comparing across trials, significantly modulated neurons were identified (Table 1). To determine whether a neuron was entrained, we computed the phase locking values (PLVu^2^)^45, 46^ of Vm to stimulation pulses and compared it to a shuffled distribution formed by randomly selecting one second long Vm across the pre-stimulation and stimulation period a thousand times (Methods). Neurons with an observed PLVu^2^ greater than the 95^th^ percentile of the shuffled distribution were deemed entrained. Using these criteria, we found that 85 of the 97 recorded neurons in both cortices were modulated, exhibiting significant change in Vm amplitude or entrainment. This result also confirms that the stimulation current threshold used was successful at evoking Vm responses in most recorded neurons.

**Table 1:** Electrical stimulation modulated various aspects of neuronal Vm.

|  |  | Activated |  |  | Suppressed |  |  | Entrained |
| --- | --- | --- | --- | --- | --- | --- | --- | --- |
|  |  | Transient | Sustained | Total | Transient | Sustained | Total |  |
| Motor Cortex | 140 Hz (n=16) | 62.5% (10) | 50.0% (8) | 81.3% (13) | 0.0% (0) | 18.8% (3) | 18.8% (3) | 37.5% (6) |
|  | 40 Hz (n=21) | 61.9% (13) | 57.1% (12) | 66.7% (14) | 4.8% (1) | 0.0% (0) | 4.8% (1) | 100.0% (21) |
| Visual Cortex | 140 Hz (n=23) | 30.4% (7) | 60.9% (14) | 65.2% (15) | 13.0% (3) | 4.3% (1) | 13.0% (3) | 87.0% (20) |
|  | 40 Hz (n=25) | 16.0% (4) | 12.0% (3) | 16.0% (4) | 56.0% (14) | 48.0% (12) | 56.0% (14) | 100.0% (25) |

Across the modulated neurons, motor cortex ones were predominantly activated, with 66.7% and 81.3% activated by 40 Hz and 140 Hz stimulation respectively (Fig. 2A, B). Interestingly, while 140 Hz in visual cortex led to a similar fraction of activated neurons (65.2%, Fig. 2C), 40 Hz primarily suppressed visual cortical neurons, suppressing 56% of the neurons (Fig. 2D). The differences in the fraction of activated and suppressed FSIs across the four testing conditions, two frequencies in two brain regions, were significant (Fisher’s test, p=1.38e^-05^, Fig. 2, Table 1). Furthermore, the time course of Vm modulation varied, with some neurons only modulated during the transient period and some only during the sustained period. Thus, while most FSIs were activated by stimulation at 40 Hz and 140 Hz, a sizable fraction of visual cortex FSIs were suppressed by 40 Hz stimulation, highlighting a brain region and stimulation frequency specific effect.

**Figure 2:**
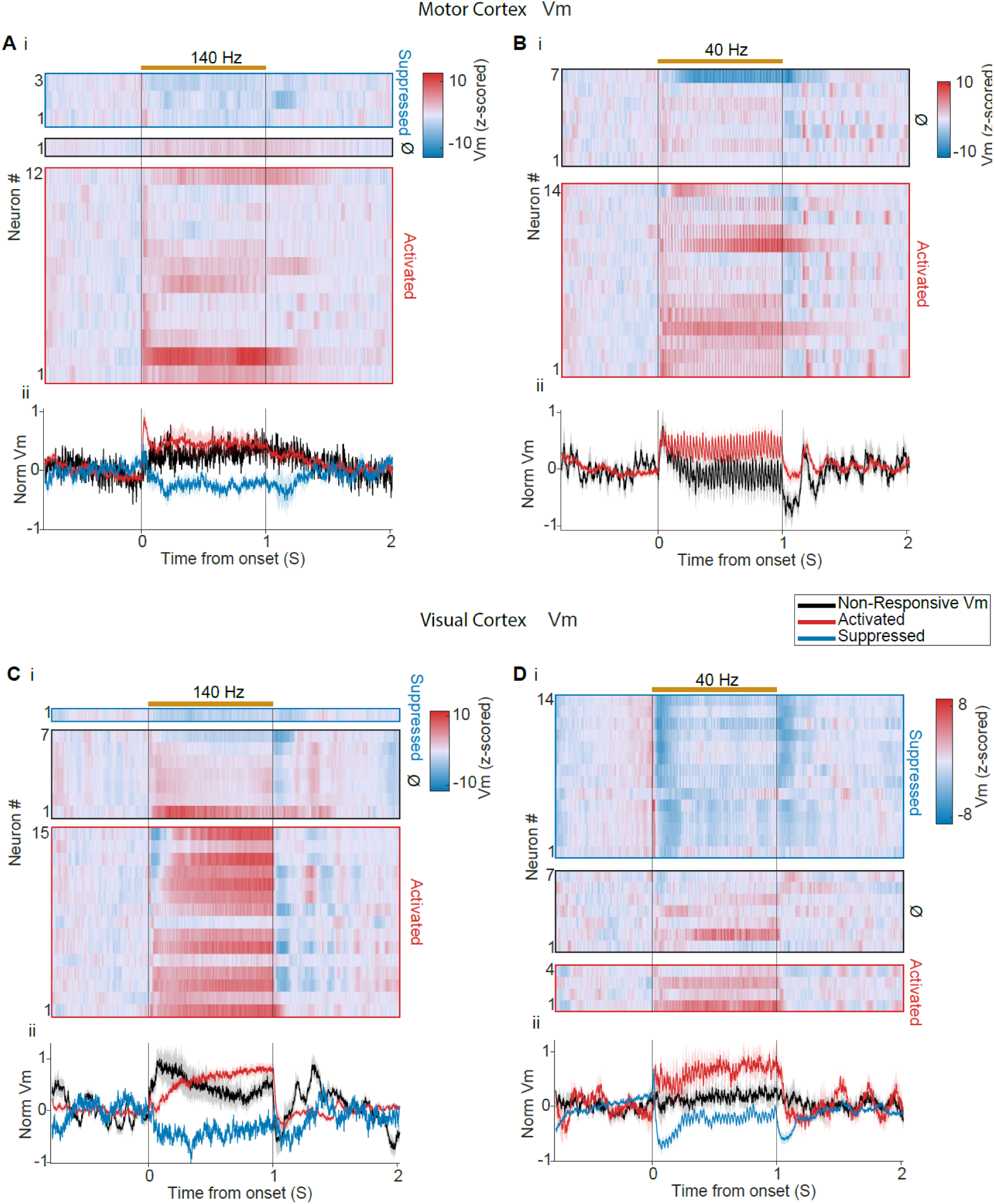
Stimulation evoked Vm depends on the stimulation frequency and brain region. (A) (i) Heatmap of evoked Vm of motor cortex FSIs upon 140 Hz stimulation, sorted into suppressed, non-modulated (Ø), or activated groups, and plotted in ascending order based on the amplitude of transient Vm changes. (ii) Population Vm across suppressed (n=3), unchanged (n=1), or activated (n=12) FSIs. (B) (i) Same as A, but upon 40 Hz stimulation. (ii) Population Vm across suppressed (n=0), unchanged (n=7), or activated (n=14) FSIs. (C) (i) Same as a, but for visual cortex FSIs upon 140 Hz stimulation. (ii) Population Vm across suppressed (n=1), unchanged (n=7), or activated (n=15) FSIs. (D) (i) Same as c, but upon 40 Hz stimulation. (ii) Population Vm across suppressed (n=14), unchanged (n=7), or activated (n=4) FSIs. Shaded areas are the standard error of the mean (SEM).

### Motor cortical FSIs are transiently and weakly entrained by 140 Hz stimulation, in contrast to the strong and persistent entrainment by 40 Hz stimulation

Previous attempts with electroencephalogram (EEG) and extracellular physiology, after removing electrical stimulation artifacts, reported some entrainment of spiking and local field potential dynamics at lower stimulation frequencies (<50 Hz)^9, 47, 48^. In our previous voltage imaging study, free of stimulation artifacts, we found that 40 Hz stimulation, but not 140Hz, robustly entrain the Vm of hippocampal pyramidal neurons^23^. Voltage imaging of motor cortical FSIs revealed that 140 Hz stimulation entrained 6 out of 16 motor cortex neurons (Table 1). Across the population of entrained neurons, Vm exhibited a rapid depolarization accompanied by a transient increase in spiking (Fig. 3A, C) and 140 Hz Vm spectral power (Fig. 3E). Aligning Vm to each stimulation pulse within the pulse train revealed that 140 Hz entrainment was stronger during the transient period with each pulse evoking a greater Vm depolarization and increased spiking than the sustained period (p=0.004, p=7.7e-6, respectively, Wilcoxon rank-sum test, Fig. 3Gii,iii, Hii,iii). Interestingly, spiking increased ∼4.8 ms after pulse onset during the transient phase, but ∼3.4 ms during the sustained phase, suggesting a circuit contribution to entrainment during persistent stimulation.

**Figure 3:**
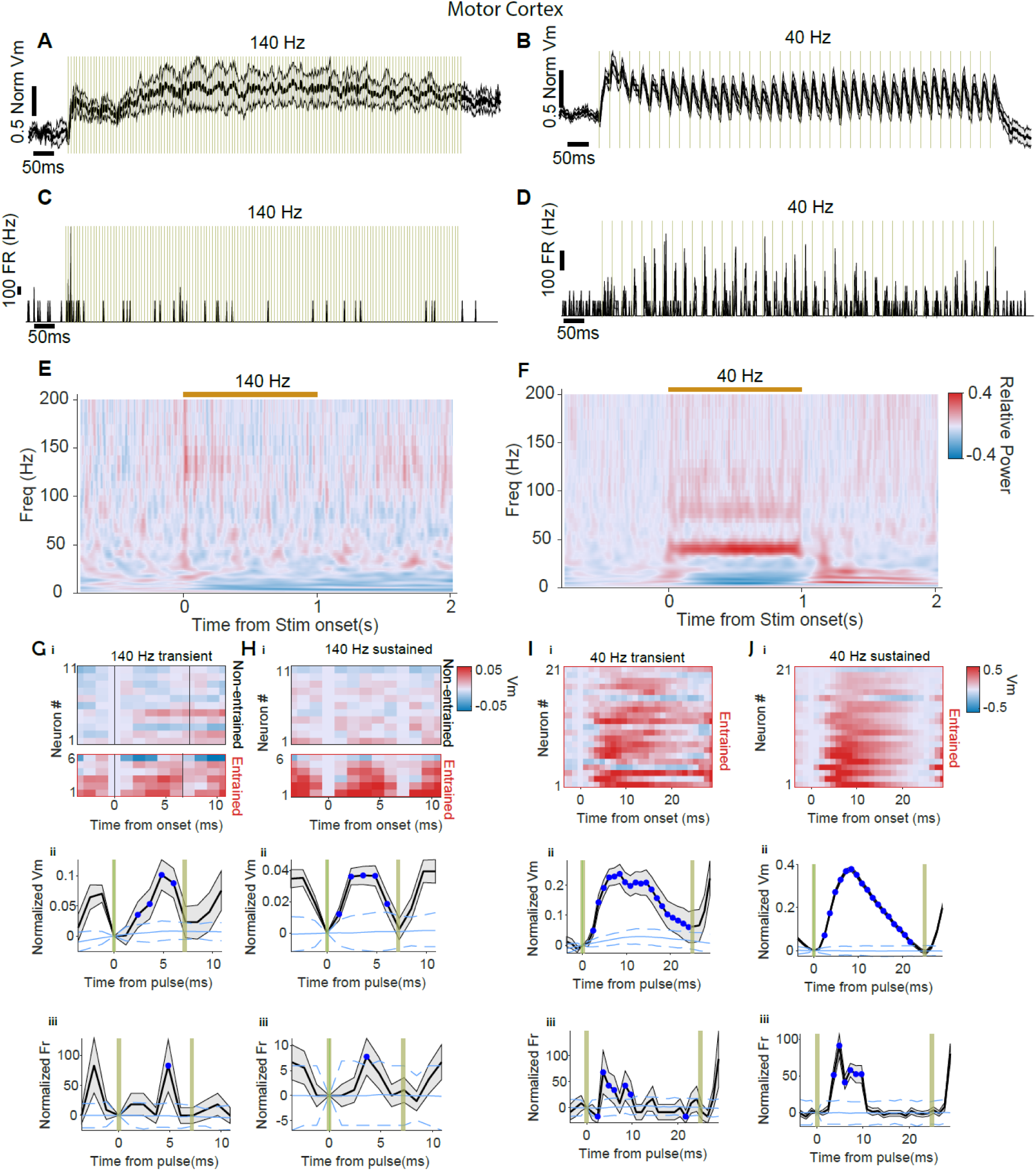
140 Hz stimulation transiently and weakly entrained Vm of motor cortical FSIs, in contrast to the strong and persistent entrainment by 40 Hz. (A) Population Vm across entrained FSIs in motor cortex upon 140 Hz stimulation. (B) Same as A, but upon 40 Hz stimulation. (C) Population firing rate across entrained neurons upon 140 Hz stimulation. (D) Same as c, but upon 140 Hz stimulation. (E) Vm spectral power across all motor cortex FSIs upon 140 Hz stimulation. (F) Same as E, but upon 40 Hz stimulation. (G) (i) Heatmap of the z-scored Vm aligned to each stimulation pulse during the transient period (100 ms) of 140 Hz stimulation onset. Neurons were grouped by entrained (n=6) and non-entrained (n=11) neurons and then sorted by increasing evoked Vm amplitude. The two black vertical lines are pulse onset and offset. (ii, iii) Pulse-triggered Vm (ii) and firing rate (iii) during the transient period. Blue line indicates the mean of the shuffled distribution, and blue dashed lines are 2.5% and 97.5% percentile of the shuffled distribution. Blue dots indicate points that are significantly different from the shuffled distribution. (H) Same as g, but for evoked responses during the sustained period. Gold vertical lines indicate individual pulses. Shaded gray area indicates SEM. (I-J) Same as g-h, respectively, but upon 40 Hz stimulation.

In contrast to the weak entrainment by 140 Hz stimulation, 40 Hz entrained all recorded motor cortex neurons, pacing both Vm and firing rate and increasing 40 Hz Vm spectral power throughout the entire stimulation periods (Fig. 3B, D, F, Table 1). Aligning Vm to stimulation pulse onset revealed a significant Vm depolarization and spiking throughout the entire 25 ms of inter-pulse interval during both the transient and the sustained periods (Fig. 3Ii-iii, Ji-iii). Spiking increase primarily occurred on the rising phase of Vm depolarization within ∼10 ms of pulse onset. Thus, 40 Hz stimulation effectively entrained motor cortex FSIs, with each stimulation pulse leading to robust Vm depolarization and spiking throughout the stimulation period.

When comparing the two stimulation frequencies, 140 Hz entrained 37.5% of motor cortex neurons, significantly lower than the 100% observed during 40 Hz stimulation (Fisher’s test, p=2.3e-5, Table 1). Since some of the non-entrained neurons could exhibit weak entrainment but not enough to pass our statistical criteria at the single cell level, we further examined the population responses across the non-entrained neurons during 140 Hz stimulation. Interestingly, across the non-entrained FSIs, stimulation pulses resulted in a small but significant Vm hyperpolarization and decrease in firing during the transient period, in sharp contrast to the Vm depolarization and increased firing detected across the entrained neurons (Fig. 3 and S4). Together, these results demonstrated that 140 Hz stimulation led to transient Vm entrainment in the motor cortex with each stimulation pulse leading to prominent Vm depolarization and spiking in entrained neurons, and weak Vm hyperpolarization and spike suppression in the non-entrained population that are most likely network driven.

### Visual cortical FSIs are robustly entrained by 140 Hz stimulation, comparable to that observed during 40 Hz stimulation

In contrast to the weak entrainment observed in the motor cortex, 140 Hz stimulation robustly entrained visual cortex neurons, with 87.0% FSIs entrained (Table 1), significantly higher than the motor cortex (Fisher’s test, p=0.002, Table 1). Across the entrained population, Vm depolarization built up gradually, leading to an overall increase in Vm 140 Hz spectral power and firing rate throughout the stimulation period (Fig. 4A, C, E). Further evaluation of Vm evoked by each stimulation pulse revealed pronounced Vm depolarization, during both the transient and sustained periods (Fig. 4Gi,ii, Hi,ii). Interestingly, while the firing rate following each pulse increased during the transient period (Fig. 4Giii), firing rate dropped during the sustained period (Fig. 4Hiii). Closer examination of the timing of the evoked Vm and spiking changes revealed that Vm depolarization peaked at ∼2.4 ms after pulse onset, whereas firing rate reduction peaked at ∼4.8 ms after pulse onset when Vm repolarized towards the pre-pulse level (Fig. 4Hii, iii). As the firing rate increased from 58±15 spikes/sec (mean ± SEM, n=20 entrained neuron) during the transient period to 90±5 spikes/sec during the sustained period, the reduction in spike rate accompanying Vm repolarization following each stimulation pulse suggests that firing rate of FSIs might have reached the upper limit during the sustained period. Distinct from that observed in the motor cortex, the few non-entrained visual cortical neurons as a population showed small but significant Vm depolarization and increased spiking following each pulse during the transient period, and no change during the sustained period (Fig. S4F, H, J).

**Figure 4:**
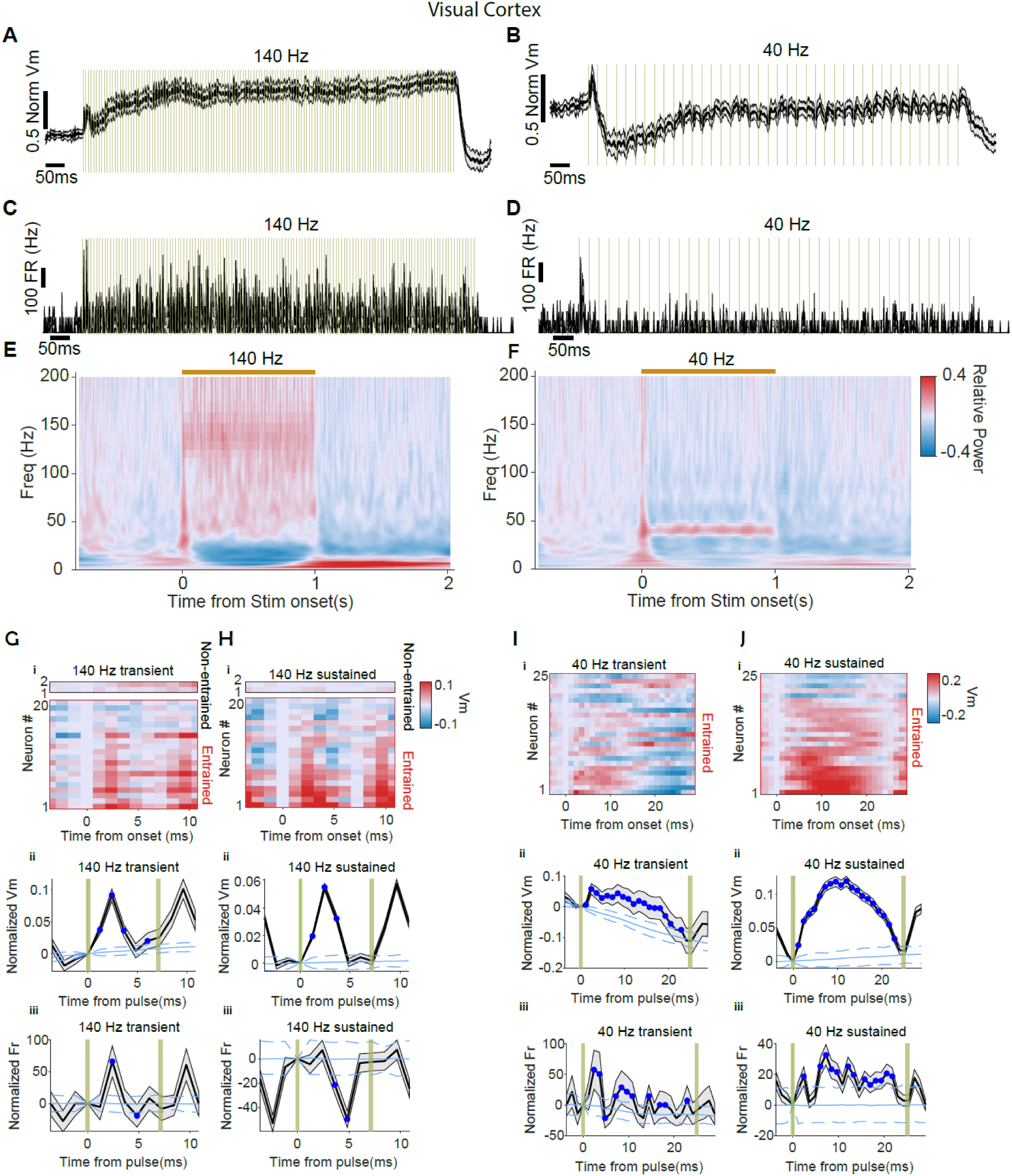
140 Hz stimulation robustly entrained Vm of visual cortical FSIs throughout the stimulation period. (A) Population Vm across entrained FSIs in visual cortex upon 140 Hz stimulation. (B) Same as a, but upon 40 Hz stimulation. (C) Population firing rate across entrained neurons upon 140 Hz stimulation. (D) Same as c, but upon 140 Hz stimulation. (E) Vm spectral power across all visual cortex FSIs upon 140 Hz stimulation. (F) Same as e, but upon 40 Hz stimulation. (G) (i) Heatmap of the z-scored Vm aligned to each stimulation pulse during the transient period of 140 Hz stimulation onset. Neurons were grouped by entrained (n=20) and non-entrained (n=2) neurons and then sorted by increasing evoked Vm amplitude. The two black vertical lines are pulse onset and offset. (ii, iii) Pulse-triggered Vm (ii) and firing rate (iii) during the transient period. Blue line indicates the mean of the shuffled distribution, and blue dashed lines are 2.5% and 97.5% percentile of the shuffled distribution. Blue dots indicate points that are significantly different from the shuffled distribution. (H) Same as g, but for evoked responses during the sustained period. Gold vertical lines indicate individual pulses. Shaded gray area indicates SEM. (I-J) Same as g-h, respectively, but upon 40 Hz stimulation.

Similar to that observed in the motor cortex, 40 Hz stimulation entrained all modulated visual cortex neurons (Table 1). Although 40 Hz stimulation predominantly evoked Vm hyperpolarization during the transient period (Fig. 4B, D), each stimulation pulse evoked a small Vm depolarization and increased spiking, which was followed by a profound hyperpolarization (Fig. 4Iii, iii), resulting in a net Vm hyperpolarization following each pulse onset. In contrast, during the sustained period, each stimulation pulse consistently led to robust Vm depolarization and spiking, which settled back to pre-stimulation levels by the subsequent onset (Fig. 4Jii, iii). Thus, while 40 Hz stimulation led to similarly robust entrainment in both motor and visual cortices, the evoked temporal Vm profiles differed, with visual cortex exhibiting a greater suppression than motor cortex, especially during the transient period.

We then further compared the entrainment effects of cortical FSIs to hippocampal CA1 pyramidal neurons reported in our previous study^23^. CA1 pyramidal neurons were robustly entrained by 40 Hz stimulation, with each pulse leading to robust Vm depolarization and spike rate increases (Fig. S5B, D). However, 140 Hz stimulation pulses led to pronounced Vm hyperpolarization and spike rate reduction (Fig. S5A, C), in sharp contrast to the overall excitatory effects observed in cortical FSIs, further highlighting cell types and brain region-dependent stimulation effects.

### Stimulation evokes distinct motor versus visual cortex FSI population responses, consistent with their different ion channel expression profiles

To further evaluate the brain region and frequency-dependent stimulation effects, we compared the population responses of all modulated neurons across conditions. In the motor cortex, 140 Hz stimulation evoked prominent Vm population depolarization and increased spiking that were restricted to the transient period (Fig. 5A, C, E, G). In contrast, 40 Hz evoked Vm depolarization persisted throughout the transient and sustained periods, though without accompanying increase in spiking (Fig. 5B, D, F, H). In the visual cortex, 140 Hz evoked significant Vm depolarization and increased spiking during both the transient and sustained periods (Fig. 5I, K, M, O), achieving a greater Vm depolarization during the sustained period than the transient period (Wilcoxon Rank Sum, p=0.00015, Fig. 5I, M). Interestingly, 40 Hz stimulation evoked a brief excitation that lasted for ∼10 ms, which was followed by a prominent inhibition lasting for ∼250 ms and returned to pre-stimulation level (Fig. 5J, N). Because of the rapid switch from excitation to inhibition, 40 Hz stimulation produced an overall Vm hyperpolarization during both the transient and the sustained period, though not significant from the baseline. But the Vm hyperpolarization is significantly less in the transient period than the sustained period (Fig. 5N), leading to a significant reduction in firing rate during the sustained period (Fig. 5L, P). The distinct effect of stimulation at 40 and 140 Hz on Vm and spiking across neuronal population in the visual versus motor cortices further underscores the brain region and frequency specific effects of intracranial stimulation.

**Figure 5:**
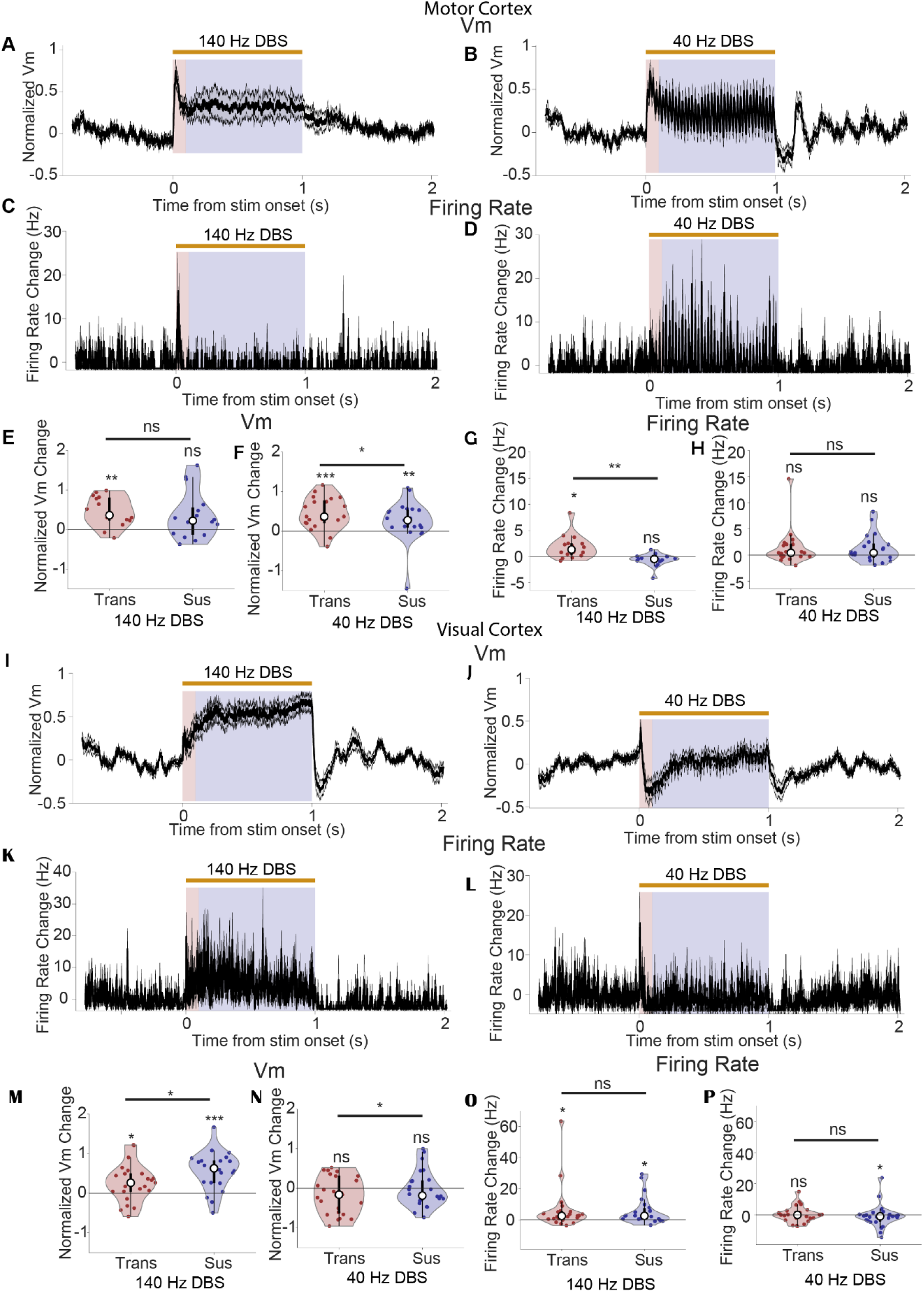
Electrical stimulation evoked population responses differ between motor and visual cortices. (A) Population Vm of motor cortex FSIs upon 140 Hz stimulation (n=16). (B) Same as A, but upon 40 Hz stimulation (n=21). (C) Population firing rate of motor cortex FSIs upon 140 Hz stimulation. Firing rates were estimated with a sliding window of 3.6 ms. (D) Same as C, but upon 40 Hz stimulation. (E) Normalized motor cortical neuron’s Vm change during the transient and sustained periods of 140 Hz stimulation. (F) Same as E, but for 40 Hz. (G) Normalized firing rate change of motor cortical FSIs during the transient and sustained periods of 140 Hz stimulation. (H) Same as g, but for 40 Hz. (I) Population Vm of visual cortex FSIs upon 140 Hz stimulation (n=23). (J) Same as I, but upon 40 Hz stimulation (n=25). (K) Population firing rate of visual cortex FSIs upon 140 Hz stimulation. (L) Same as k, but for 40 Hz. (M) Normalized visual cortical neuron’s Vm change during the transient and sustained periods of 140 Hz stimulation. (N) Same as m, but for 40 Hz stimulation. (O) Normalized firing rate change of visual cortical FSIs during the transient and sustained periods of 140 Hz stimulation. (P) Same as O, but for 40 Hz stimulation. Shaded gray area is the SEM. Shaded red area indicates the transient period. Shaded blue area indicates the sustained period. Gold bar indicates stimulation period. (Sign Test and Wilcoxon’s Signed Rank Test, statistical results in **Ext. Data Table 1**, ns = non-significant, **<0.01, and ***<0.001).

To further explore the cellular properties that may contribute to the distinct stimulation-evoked population responses observed across frequencies and brain regions, we analyzed the gene expression profiles of common ion channels and pumps using the Allen Institute single-cell transcriptomes data (https://portal.brain-map.org/atlases-and-data/rnaseq/mouse-whole-cortex-and-hippocampus-10x) that contains 1.1 million cortical cells^43, 49, 50^. We found that parvalbumin positive FSIs in the visual and motor cortices have comparable levels of many channels and pumps relevant to neuronal excitability, including various sodium, potassium, and calcium channels, and Na/K-ATPase (Fig. S6A-C). Some channels, e.g. Kcna1, Kcnc3, Cacna1d, and Hcn1, exhibited negligible differences between motor and visual cortices (Wald test, p>0.05, cliff’s delta < 0.147, Fig. S6C-O). However, Cacna1a (encoding Cav2.1 calcium channel) had slightly higher expression in the visual cortex (Wald test, p=0.0, cliff’s delta=0.18, Fig. S6I), whereas Kcnc1 (encoding Kv3.1) (Wald test, p=0.0033, cliff’s delta=0.18, Fig. S6E) and Kcna2 (encoding Kv1.2) (Wald test, p=0.0033, cliff’s delta=0.18, Fig. S6E) had higher expression in the motor cortex. These differences highlight the variation in biophysical properties among FSIs between the two cortical regions, which may contribute to the distinct population Vm responses observed during stimulation.

### Electrical stimulation reduces FSI response amplitude to visual flicker inputs and bidirectionally impacts response timing

Electrical stimulation provides an external input to a neuron and thus influences neuronal information processing^22, 23^. To evaluate how electrical stimulation evoked response influences neuronal processing of intrinsic synaptic inputs, we characterized the effect of electrical stimulation on visual cortical FSIs upon visual flicker presentation. Specifically, we performed voltage imaging of FSIs while presenting 8 Hz visual flickers delivered via a white LED to awake mice (Fig. 6A). Each trial lasted for 5 seconds, consisting of a 1 second baseline, 3 seconds of visual flickers, and 1 second post-flicker. Electrical stimulation at 40 Hz or 140 Hz was delivered for 1 second in the middle of the visual flicker presentation period (Methods, Fig. 6B).

**Figure 6:**
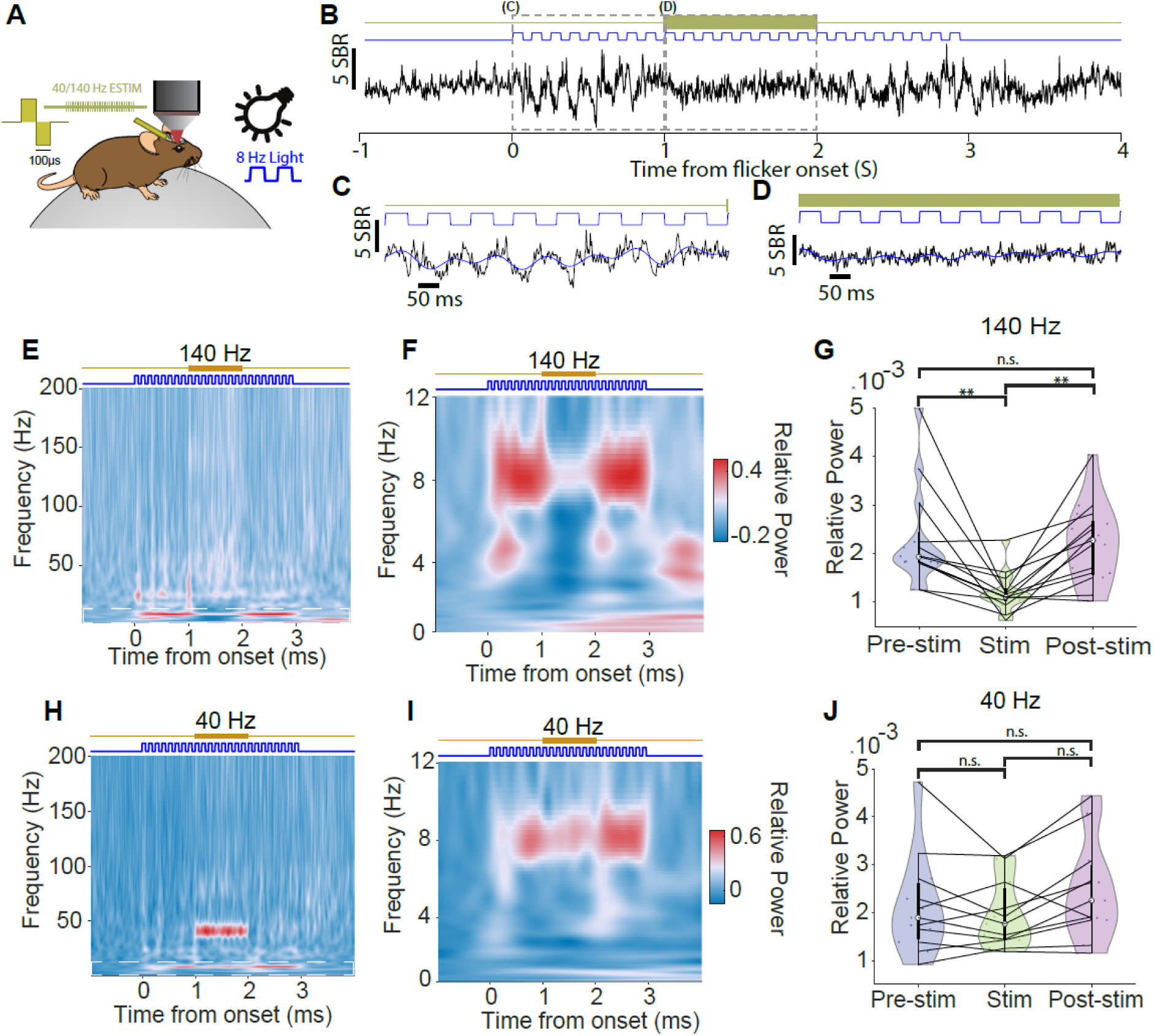
140 Hz stimulation, but not 40 Hz, disrupts FSI cellular processing of 8 Hz visual flicker inputs. (A) Experimental setup illustrating simultaneous voltage imaging, visual flicker presentation, and electrical stimulation. (B) Example FSI Vm responses throughout an experimental trial. (C) Zoom in view of Vm during the 1 second with visual flicker presentation (0-1 second in b). (D) Zoom in view during simultaneous visual flicker and 140 Hz stimulation (1-2 seconds in b). (E) Population Vm power spectra aligned to visual flicker onset upon 140 Hz stimulation (n=14). (F) Zoom in view of the power spectra at 0-12 Hz in e. (G) Visual-flicker evoked 8 Hz Vm spectral power before electrical stimulation (Pre-stim), during 140 Hz stimulation (Stim), and after electrical stimulation (Post-stim). (H-J) Same as e-g, but with 40 Hz stimulation. (Kruskal-Wallis Test and Post-hoc Dunn Test, statistical results in **Ext. Data Table 2**, ns = non-significant, **<0.01, and ***<0.001).

We found that 8 Hz visual flickers alone reliably entrained Vm (Fig. 6B, C, S7), increasing the 8 Hz Vm spectral power (Fig. 6E-F, H-I). Upon electrical stimulation at 140 Hz, many FSIs exhibited a reduced Vm response to flickers (Fig. 6D, E-F, H-I). Of the 13 neurons tested with 140 Hz electrical stimulation, 6 (46%) showed a significant reduction in flicker-evoked 8 Hz Vm spectral power, which recovered at electrical stimulation offset (Fig. 6G). In contrast, of the 11 neurons tested with 40 Hz stimulation, only 1 (9%) showed a significant reduction in flicker-evoked 8 Hz spectral power, and 40 Hz stimulation failed to alter the population responses to flicker-evoked responses (Fig. 6J). Thus, electrical stimulation at 140 Hz, but not 40 Hz, effectively attenuated the response amplitude of FSIs to visual flickers.

**Figure 7:**
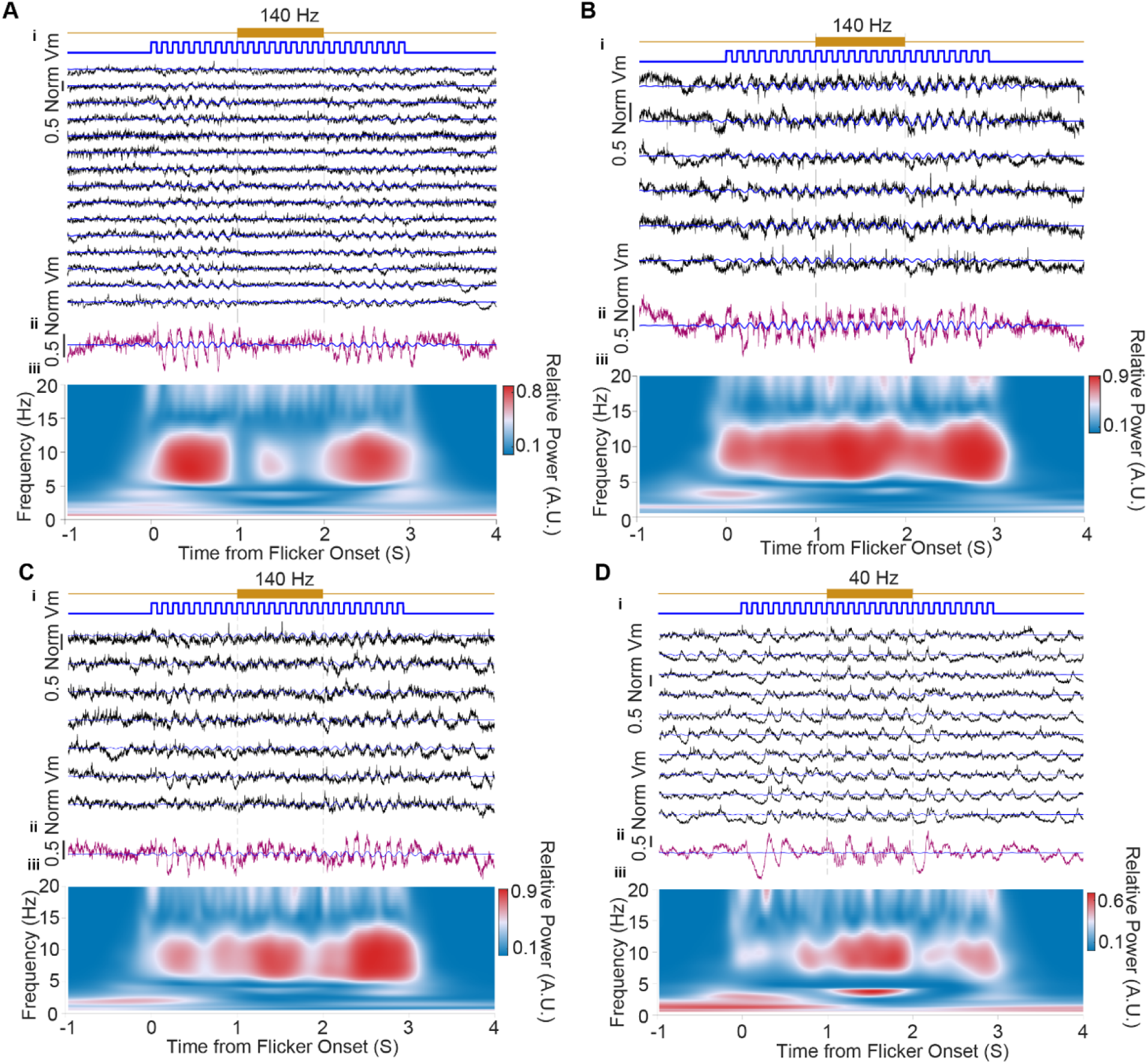
Electrical stimulation bidirectionally modulates the temporal precision of visual cortical FSI response to visual flickers. A) An example FSI showing decreased 8 Hz coherence between its Vm and flicker timing during 140 Hz stimulation. (i) Vm traces across experimental trials. (ii) Trial-averaged Vm. (iii) Trial-averaged coherence across frequencies between Vm and visual flickers. B) Same as a, but for a FSI showing increased 8Hz coherence during 140 Hz stimulation. C) Same as b, but a different FSI. D) Same as c, but for a FSI showing increased 8 Hz coherence during 40 Hz stimulation.

To further assess how electrical stimulation influences FSI response timing to individual flickers within the flicker train, we quantified the coherence between Vm traces and flicker timing. We found that electrical stimulation reduced the coherence only in 1 neuron, which was during 140 Hz stimulation, highlighting that electrical stimulation could disrupt the response timing to visual inputs (Fig. 7A). Intriguingly, stimulation increased coherence in 3 neurons, 2 during 140 Hz (Fig. 7B-C) and 1 during 40 Hz (Fig. 7D), suggesting that electrical stimulation could enhance the temporal precision of visual information processing. Together, these results demonstrate that electrical stimulation at either 40 Hz or 140 Hz modulates visual cortex FSI response to natural sensory inputs. While electrical stimulation generally reduces the response amplitude to sensory inputs as measured by reduced Vm spectral power, it exerts diverse effects on the temporal precision of visual cortical FSI processing of sensory inputs.

## Methods

### Animal Preparation

All animal experiments were performed in accordance with the National Institute of Health Guide for Laboratory Animals and approved by the Boston University Institutional Animal Care and Use and Biosafety Committees. 11 adult PV-Cre mice (JAX, 129P2-Pvalbtm1(cre) Arbr/J, 8 Male and 3 Female), 8-16 weeks old at the start of the experiments, were used, with 6 (5 male and 1 female) for motor cortex experiments and 6 (4 male and 2 female) for visual cortex experiments. No statistical method was used to predetermine the sample size of mice groups.

Under 0.5-2% isoflurane aesthesia, we first created a 3 mm diameter craniotomy over the motor cortex (centered on AP: +1.5 mm, ML:-1.5 mm) or the visual cortex (centered on AP:-2.8 mm, ML: +2.5 mm). We then infused 500 μl AAV-CAG-FLEX-SomArchon-GFP (University of North Carolina vector core, titer: 6.3e12 gc/mL) at a flow rate of 100 nL/min at 3 locations within each craniotomy using an ultra-micropump infuser (World Precision Instruments, UMP3) with the tip terminating at ∼250 μm below the cortical surface. An electrode made of insulated stainless-steel wire (PlasticsOne, 005SW/30S,) soldered to a 0.016” diameter dip pin (Mill-Max, series 853-93, ED90571-ND) was inserted into the brain tissue anterior to the craniotomy. Another electrode was inserted posterior to the craniotomy to serve as the ground for electrical stimulation. We then placed a circular coverslip (Deckgläser Cover Glasses, Warner Instruments Inc., 64-0726, #0, OD: 3 mm) over the craniotomy and sealed to the surrounding bone via UV curable glue (Norland Products Inc., Norland Optical Adhesive 60, P/N 6001). The coverslip and the electrodes were secured onto the skull using Metabond (C&B-Metabond® Clear L-Powder). Dental cement (Stoeltingco #51459 Clear) was applied to further secure the coverslip, electrodes, and a custom aluminum bar to the skull. Sustained release buprenorphine (Reckitt Benckiser Healthcare; 0.03 mg/kg, i.m.) was administered preoperatively to provide continued analgesia for >72 hours.

### SomArchon Voltage Imaging

We first habituated mice to head fixation and a pinned spherical Styrofoam ball as previously described in Shroff et al^44^ SomArchon imaging was conducted with a custom widefield microscope, as in Shroff et al^44^, or a custom targeted illumination confocal (TICO) microscope, as detailed in Xiao et al^65^. The widefield microscope was equipped with an ORCA-Fusion Digital complementary metal oxide semiconductor (CMOS) Camera (Hamamatsu Photonics K.K., C14440-20UP), a 40X NA=0.8 water immersion objective (Nikon), a 140 mW 637 nm red laser (Coherent, Obis 637-140), a 635 nm dichroic filter (Semrock, FF635-Di01-25×36), and a 664 nm long pass emission filter (Olympus, OCT49006BX3). A mechanical shutter (Newport corp., model 76995) was positioned in the laser path to control the timing of SomArchon excitation via a NI DAQ board (USB-6259, National instruments). SomArchon fluorescence was acquired at 500 or 833 Hz using the HCImage Live software (Hamamatsu Photonics), and recorded files were stored as DCAM imaging files (DCIMG) and analyzed offline with MATLAB (Mathworks Inc.). The TICO microscope was equipped with a sCMOS camera (Teledyne Photometric, Kinetix), a 16X (NA=0.8, LWD, Nikon) objective, a 637 nm red laser (Ushio America Inc., Red-HP-63X), two dichromatic mirrors (Thorlabs DMSP550R, DM4; Thlorlabs DMLP605R, DM5), and a quadband dichromatic mirror (DM1, Chroma Technology Corp. ZT405/488/561/640rpcv2). A mechanical shutter (Thorlabs, SC10) controlled the timing of SomArchon excitation via a NI DAQ board (National instruments, USB-6259). SomArchon fluorescence was acquired at 800 Hz using the PmTestSuite software, and the recorded files were stored as RAW format image files and analyzed offline with MATLAB.

### Electrical Stimulation

Electrical stimulation was delivered at 40 Hz or 140 Hz for 1 second, during the middle of each 3-second-long voltage imaging trial, using an isolated pulse stimulator (A-M Systems Model, 4100). Inter-trial-interval was 10 seconds. Electrical stimulation pulses were biphasic, square-wave, and charge balanced, at 200 μs per phase. Due to the variability across animals (e.g. variation in local tissue architecture, electrode impedances and positions), we first empirically determined the current threshold for each animal. Specifically, we delivered 40 Hz stimulation, starting at 10 μA with 10 μA increments, until the current amplitude that evoked any change in SomArchon fluorescence amplitude or spectra power. This current was then deemed the minimal current for the given animal. During each subsequent imaging session, we further tuned the current, starting at the minimal current, and increased with 10 μA increments until a change in fluorescence of the recorded neuron was detected. After determining the current threshold, neurons were recorded for 10-15 trials, with each trial lasting 3 seconds with a 10-second inter-trial-interval. Occasionally, we encountered neurons that did not respond to current at 200% of the minimal current, and these neurons were not investigated further. The stimulation pulse and SomArchon image frame timestamps were recorded by OmniPlex (PLEXON, Omniplex) or Open Ephys (Open Ephys, SKU: OEPS-9030), and aligned offline.

### Electrical stimulation during visual flicker presentation

To test the effect of electrical stimulation on visual flicker evoked neuronal changes, each neuron was recorded for 10 trials. Each trial contained a 5 second SomArchon recording with 1 second baseline, 3 second visual flicker presentation, and 1 second post flicker. The inter-trial-interval was ∼30 seconds. 1 second of electrical stimulation was delivered in the middle of the visual flicker presentation. Electrical stimulation pulses were biphasic, square-wave, and charge balanced, at 100 μs per phase. Visual flickers were presented via a 5 mm white LED (Amazon, MCIGICM) at 8 Hz, 50% duty cycle, at an intensity of 20.5 μW in a dimly lit room. LED was controlled via TTL pulses generated through a DAQ (National Instruments, BNC-2110 and PCIe-6323). The LED on and off timestamps were recorded by Open Ephys (Open Ephys, SKU: OEPS-9030) at an acquisition rate of 30 kHz, and aligned to SomArchon image frames offline.

### SomArchon Trace Pre-processing and Spike identification

SomArchon fluorescence image frames were first motion corrected and segmented as detailed previously^23, 66^. The SomArchon trace for each neuron was then extracted, and detrended by subtracting the exponential fit of the fluorescence before and after stimulation, ensuring that stimulation-evoked changes did not influence the fit. The resulting traces were then considered the membrane voltage (Vm) traces for each neuron. All Vm traces across individual trials were manually inspected to discard any trial that exhibited unstable fluorescence. We then detected spikes from the Vm traces as that described previously^67^. Briefly, we first high-pass filtered (>120 Hz) the trace to remove slow changes and then computed an “upper trace” and an “lower trace” of the filtered Vm to separate fast spike-related activity from slow fluorescence fluctuations. To generate the “upper trace” and “lower trace”, we computed a “smoothed trace” by smoothing the trace with a moving window average of ± 500 frames. Fluorescence points in the original trace that were above the “smoothed trace” were assigned to the “upper trace”, while fluorescence points below the “smoothed trace” were assigned to the “lower trace”. Spikes were identified as fluctuations in the “upper trace” that were greater than a threshold, set as 3.75 times the standard deviation of the fluctuations in the “lower trace”.

To estimate the spike to baseline ratio (spike SBR), spike amplitude was calculated by taking the difference between the height of the detected spike and the lowest of its neighboring points. To calculate the baseline noise, we first computed a “spikes-removed trace” by removing the detected spike peaks and the surrounding three points before and after spikes. Spike SBR was then computed by dividing the spike amplitude by standard deviation of the “spikes-removed trace.” Spike rate was computed using a moving window of 3.6 ms for Fig. 5 C, D, K, L.

### Classifying Vm and Firing Rate Modulation of Single Neurons

To determine whether electrical stimulation evoked significant changes in a neuron’s Vm, we compared the Vm during the transient (0-100 ms of onset) or sustained period (100-1000 ms of onset) to the corresponding period before stimulation onset (-500 to 0 ms) using the Wilcoxon sign-rank test (p<0.05). Neurons with a significantly higher Vm after stimulation onset than before the onset were classified as activated. Neurons with a significantly lower Vm after onset than before the onset were classified as suppressed.

### Vm power spectra density analysis

Vm spectral power was estimated using a continuous wavelet transform with the Morse wavelet function, implemented in MATLAB’s *cwt*, across a frequency range of 1-200 Hz. Each frequency component across time was computed as the absolute value of the complex numbers. For the 3 second trials with 1 second of stimulation, the power was normalized by the following formula:

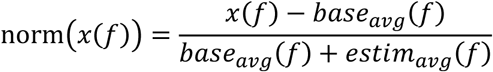

*base_avg_(f)* was the average power in the 1 second baseline period before stimulation onset at frequency *f*. *estim_avg_(f)* was the average power during the stimulation period at frequency *f*. *x(f)* is the instantaneous power at frequency *f*.

For the 5 second simultaneous visual flicker and electrical stimulation trials, the power was normalized by the following formula:

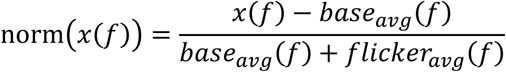

The *base_avg_(f)* was the average power in the 1 second baseline period before visual flicker onset. The *flicker_avg_* was the 1 second period at the onset of visual flicker before electrical stimulation onset. Power spectra for each neuron were calculated for each trial and then averaged across all trials to get a neuron’s trial-averaged power spectra.

### Coherence between neuronal Vm to visual flicker timing

Coherence was calculated using MATLAB’s *wcoherence* function over a frequency range of 0-200 Hz. For each trial, coherence was computed by inputting the Vm trace and the visual flicker raster recorded by the OpenEphys system. The final coherence value for each neuron was then computed by averaging the trial-wise coherence across all trials.

### Phase locking analysis

We first estimate the consistency of Vm phase relative to the timing of each stimulation pulse by computing phase-locking value (PLV)^46^:

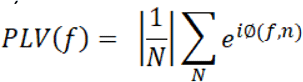

where *f* is the Vm frequency and *N* is the total number of events. The phase *ϕ* of the Vm at pulse time was obtained from the Hilbert Transform of the narrowband filtered Vm, ±5% of *f*. PLV(*f)* was computed across the frequency range of 1-200 Hz.

To avoid potential bias in the PLV due to low spike counts, only neurons with more than 10 spikes were included. Further, the spike PLV value was adjusted by spike counts using the following equation to obtain the adjusted PLV (PLV u^2^) as in a previous study^45^:

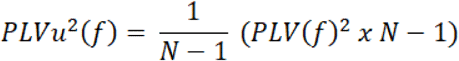

To determine whether a neuron was entrained by stimulation, we performed a permutation test comparing PLVu^2^ during stimulation versus a shuffled distribution. To form the shuffled distribution, we computed the PLVu^2^ of a randomly chosen 1 second-long continuous time window within the baseline and stimulation periods for each shuffle. This process was repeated for 1,000x to form the shuffled distribution. A neuron was classified as entrained if its observed PLVu^2^ exceeded the 95^th^ percentile of the shuffled distribution.

### Characterization of individual electrical pulse evoked Vm and firing rate changes

To determine whether individual stimulation pulses evoked significant Vm changes, we first normalized pulse-triggered Vm by subtracting the Vm at each pulse onset and then computed the mean across all pulses in all neurons. We then compared this population curve to a shuffled distribution. To generate the shuffled distribution, we averaged randomly chosen windows (having same size number of trace points within a pulse window of the respective stimulation frequency) across the stimulation period in all neurons. This was repeated 1,000x. Vm >97.5% or <2.5% of the shuffled distribution were deemed significantly modulated. Individual stimulation pulse evoked firing rate changes were identified similarly.

### Histology

Mice were perfused with phosphate-buffered saline (PBS) containing 4% paraformaldehyde. Brains were then extracted, post-fixed in 4% paraformaldehyde for up to 4 hours, and placed in 30% sucrose solution at 4°C. Once the brain was completely submerged, it was sliced on a frozen microtome (Leica, SM2010R) at 50 μm thickness. To stain for parvalbumin, slices were incubated in PBS containing 0.5% Triton (SigmaAldrich, 9036-19-5) and 100 mM glycine (SigmaAldrich, 56-40-6) for 30 minutes, followed by incubation in a blocking solution containing 2% goat serum (Jackson Laboratories ImmunoResearch 005-000-001) for 2 hours. Slices were then stained with the rabbit PV27 antibody (SWANT PV27,1:500) overnight at 4°C, followed by goat anti-rabbit with conjugated Alex Fluor 568 (Invitrogen A11011, 1:500) for two hours at room temperature. All images of antibody-stained slices were taken using an Olympus FV3000 scanning confocal microscope using a 20X objective (LUCPLFN20X; Olympus), and analyzed with *Fiji,* an open-source image processing software based on ImageJ^68^.

### Allen brain institute gene expression analysis

Single cell RNA-seq data was obtained from the Allen Brain Institute database, https://portal.brain-map.org/atlases-and-data/rnaseq/mouse-whole-cortex-and-hippocampus-10x^49^. Parvalbumin positive cells were identified as ‘Pvalb’ clusters. Parvalbumin positive cells in primary motor cortex were identified by the label ‘MOp’, and parvalbumin positive cells in primary visual cortex were identified by the label ‘VISp’. Gene expression for each Pvalb cluster was computed as the mean across neurons with 10-90^th^ percentile of the expression counts. Counts were normalized for each cell by dividing the total counts of all genes for that given cell, multiplying by 100,000, adding 1, and then performing a log2 transform as detailed in Tasic et al. and Sherman et al. ^43, 50^.

Differential gene expression was determined with the python package, https://pydeseq2.readthedocs.io/en/latest/index.html^69^. For the individual ion channel genes, p-values were determined by the Wald test as implemented in the *pydeseq2* package. Size effect was determined by calculating the cliff’s delta (δ) between parvalbumin positive cells in motor cortex and visual cortex, |δ| < 0.147 considered negligible, 0.147 ≤ |δ| < 0.330 considered small, 0.330 ≤ |δ| < 0.474 considered medium, and |δ| ≥ 0.474 considered large.

## Discussion

We characterized the effect of high frequency intracranial stimulation on FSIs in the awake mammalian brain through single cell voltage imaging of motor and visual cortical parvalbumin expressing neurons in head-fixed mice. At intensities sufficient to evoke consistent cellular Vm changes without influencing behaviors, local stimulation at 40 Hz and 140 Hz led to diverse Vm changes across neurons. While largely excitatory effects were observed during stimulation across frequencies and brain regions, visual cortical FSIs were powerfully inhibited by 40 Hz stimulation. Furthermore, visual cortical FSIs were robustly entrained throughout 140 Hz stimulation, comparable to that observed during 40 Hz stimulation. In contrast, motor cortex FSIs and hippocampal pyramidal cells, though robustly entrained by 40 Hz, were weakly entrained by 140 Hz stimulation and only during the transient 100 ms of stimulation onset. The observed brain region specific responses may be partially attributed to differences in common ion channel expression profiles, in addition to circuit variations. Finally, we found that the response amplitude of visual cortex FSIs to visual flickers was consistently attenuated by electrical stimulation, especially by 140 Hz. However, the response timing was bidirectionally modulated with some showing decreased temporal precision, while others increased temporal precision. Together, these results demonstrate brain region-and stimulation frequency-specific effects of electrical neuromodulation, with FSIs in the visual cortex, but not the motor cortex, faithfully following high frequency 140 Hz stimulation and exhibiting powerful inhibition during 40 Hz stimulation. These effects though consistently attenuate the amplitude of FSI responses to sensory inputs, bidirectionally modulate the temporal precision of cellular responses to inputs, highlighting the potential of selectively enhancing or reducing neuronal informational processing through tuning stimulation parameters.

### Evoked responses evolved throughout the stimulation period

Using voltage imaging, we recorded the real time effect of electrical stimulation, free of electrical stimulation artifacts. We found that evoked Vm responses evolved rapidly after stimulation onset, switching between depolarization and hyperpolarization within tens of milliseconds (Fig. 5). Brief bursts of electrical stimulation on the order of a few seconds is commonly used in closed-loop Responsive Neurostimulation (RNS) system for drug-resistant epilepsy^51, 52^. As epilepsy is often characterized by overexcitation^53, 54^, selective activation of inhibitory neurons via optogenetic and chemogenetic methods led to long lasting reduction of seizure risk^51, 55, 56^. The observed evolution of Vm dynamics during the brief one-second window suggests that FSIs may exhibit diverse response dynamics under chronic stimulation, and augmenting feed-forward inhibition via elevated FSI activation could be advantageous in producing network inhibition. Clinically, chronic stimulation is employed in the treatment of essential tremor, Parkinson’s disease, and various other neurological and psychiatric diseases with therapeutic benefits for different symptoms emerging over a wide time course, from seconds to hours and days^2, 57^. The delayed effects may reflect longer term synaptic plasticity induced by chronic stimulation^58^. Given that some FSIs can be entrained and activated at 140 Hz (Figs. 2A,C; 3G, H; 4G, H), prolonged activation of FSIs could lead to sustained suppression of cortical networks, shaping network plasticity.

### Brain region-dependent responses to electrical stimulation

FSIs in motor and visual cortices exhibit drastically distinct responses to local intracranial stimulation. During 40 Hz stimulation, visual cortex FSIs were robustly suppressed (Fig. 2D, Table 1), whereas motor cortex FSIs were activated (Fig. 2B). Additionally, during 140 Hz stimulation, visual cortex FSIs were consistently entrained (Fig. 4E, G, H), whereas motor cortex FSIs showed little entrainment beyond the initial 100 ms of stimulation onset (Fig. 3E, G, H). While FSIs in both cortices exhibited a sharp Vm depolarization at stimulation onset, peaked within about 20 ms (Fig. 5), CA1 pyramidal cells Vm gradually ramped and peaked well beyond 100 ms of stimulation onset^23^. Many factors could have contributed to this region-specific effect, including ion channel expression profiles as we demonstrated, and other circuit variations.

Of the various common ion channels examined, motor cortex has a greater expression level of several potassium channels (Kcna2, Kcna6, Kcnc1, Kcnab1), kainite receptors (Grik1), Na/K transporter (ATP1a1) and L-type Ca²⁺ channels (*Cacna1c*). In contrast, visual cortex has a greater level of N-type (Cacna1b) and P/Q-type Ca²⁺ channels (*Cacna1a*), BK channels (Kcnma1), potassium channels (Hcn2), and NMDA receptors (Grin1) (Fig. S6). It is difficult to directly associate the contribution of each ion channel to the observed response variations between these brain regions. Moreover, the PV population itself is heterogeneous, comprising morphologically and functionally distinct subtypes, notably basket and chandelier neurons^39^. Such structural and functional diversity could influence the response of each subtype to electrical stimulation^59^. Future modeling studies combined with causal manipulation of ion channels could reveal the biophysical mechanisms underlying their response variations to high frequency stimulation.

Stimulation-evoked activation of FSIs will lead to lateral inhibition of other connected FSIs and pyramidal cells, and thus the evoked responses detected reflect a balance between direct stimulation effect and indirect network effect. It is highly plausible that the predominantly inhibitory effects observed during 40 Hz stimulation reflect lateral inhibition between neighboring FSIs. Indeed, FSIs are known to promote network gamma oscillations^60^, and thus they could be preferentially recruited by 40 Hz stimulation. Particularly, the difference in the timing of peak Vm depolarization and spike increase following each pulse during the transient and sustained period of 140Hz stimulation in both cortices underscores the contribution of network inputs in shaping the evoked responses (Fig. 3G, H, and 4G, H). However, it is also possible that stimulation leads to short-term synaptic depression, e.g. synaptic vesicle depletion^61^ and postsynaptic receptor desensitization^61, 62^.

### Robust entrainment of visual cortical FSIs by 140 Hz stimulation

It is generally thought that Vm dynamics are not fast enough to track each individual stimulation pulses within high frequency pulse trains beyond a hundred hertz. We observed that visual cortical FSIs were robustly entrained by 140 Hz stimulation (Fig. 4E, G, H), suggesting that higher frequencies of several hundred hertz may be more effective at creating informational lesions in these neurons^63, 64^. In contrast to the robust entrainment effect observed in visual cortical FSIs, hippocampal pyramidal cells and motor cortex FSIs were poorly entrained by 140 Hz stimulation. While the lack of entrainment of hippocampal pyramidal cells is consistent with pyramidal cells having lower firing rates^27^, the failure of entraining motor cortex FSIs again reflect brain region differences.

### Electrical stimulation bidirectionally modulates the temporal precision of cellular information processing

The functional informational lesion hypothesis poses that electrical stimulation disrupts cellular processing of synaptic inputs within the targeted nucleus^22–25^, therefore blocking the propagation of pathological dynamics across interconnected circuits. We found that 140 Hz, but not 40 Hz, consistently reduced the response amplitude of FSIs to visual flickers presented to awake mice, measured as Vm spectral power (Fig. 6). This reduction in response amplitude is consistent with that observed previously in hippocampal CA1 during artificial optogenetic stimulation^23^, and the general observation that higher frequency stimulation is more therapeutic than lower frequency stimulation.

However, we also found that the temporal precision of visual flicker-mediated Vm changes, measured as the coherence between Vm and visual flickers, were bidirectionally modulated by electrical stimulation, with some neurons showing reduced temporal precision, while others showing enhanced precision (Fig. 7). One possibility is that electrical stimulation increases membrane conductance which reduces the amplitude of the evoked Vm responses and bias neuronal responses to larger synaptic inputs. Essentially, weaker synaptic inputs would fail to produce spiking output and thus increase the signal-to-noise ratio during cellular input-output information transformation. Future experiments that directly measure membrane conductance or simultaneously record multiple FSIs could help provide mechanistic insights. Nonetheless, the observed bidirectional effect on temporal processing of synaptic inputs, combined with consistent response amplitude reduction, demonstrates that electrical stimulation reshapes information coding ability of the targeted network.

## Acknowledgments

We thank members of Han Lab for their help throughout the study. X. H. acknowledges funding from the NIH (1RF1NS129520, 1R01NS119483, 1R01MH122971, and 1R01NS115797) and NSF (2002971-DIOS, and 1955981-CIF).

## Author contributions

P.F., E.L collected and analyzed the data. K.K., Y.W., Y. Z. provided technical help. P.F. and X.H. prepared the manuscript, and all authors edited the manuscript. X.H. oversaw all aspects of the project and supervised the study.

