## Supplemental Material for "High frequency electrical stimulation entrains fast spiking interneurons and bidirectionally modulates information processing"

### Supplemental Figures

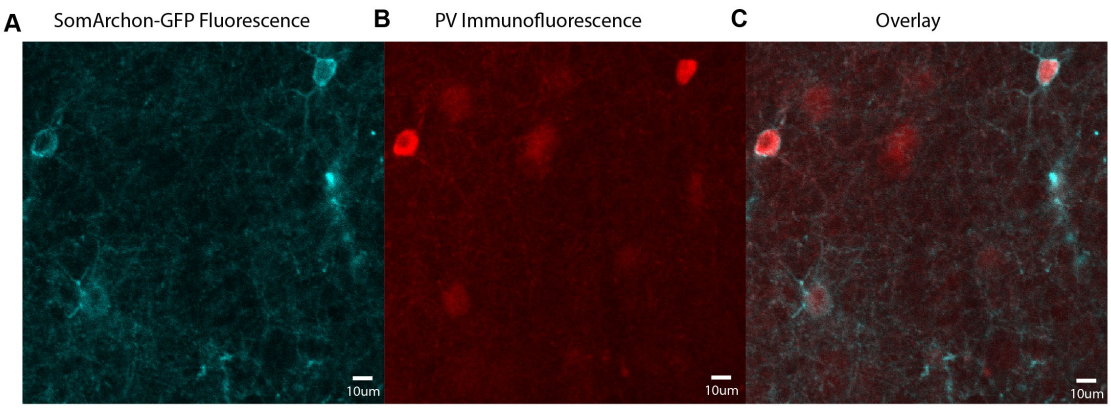


***Figure S1: SomArchon-GFP fluorescence and parvalbumin (PV) immunofluorescence in cortical FSIs****. (A) An example brain slice showing motor cortex neurons expressing SomArchon-GFP. (B) PV immunofluorescence. (C) Overlay of a and b.*

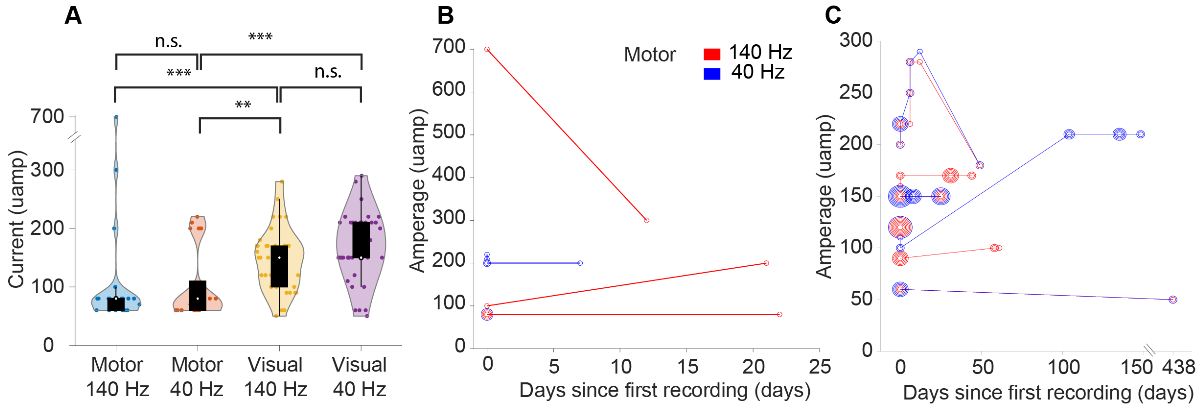


***Figure S2: Current amplitudes for recordings in the motor and visual cortices.*** *(A) The current amplitude used for each recording in the motor and visual cortex during 140 Hz and 40 Hz stimulations. (B) The current amplitude used for each neuron in the motor cortex vs the days from the first recording day. Marker size corresponds to the number of neurons recorded at the same time point with the same amplitude. Lines connect neurons recorded from the same mouse. (C) Same as b, but for visual cortex neurons.*

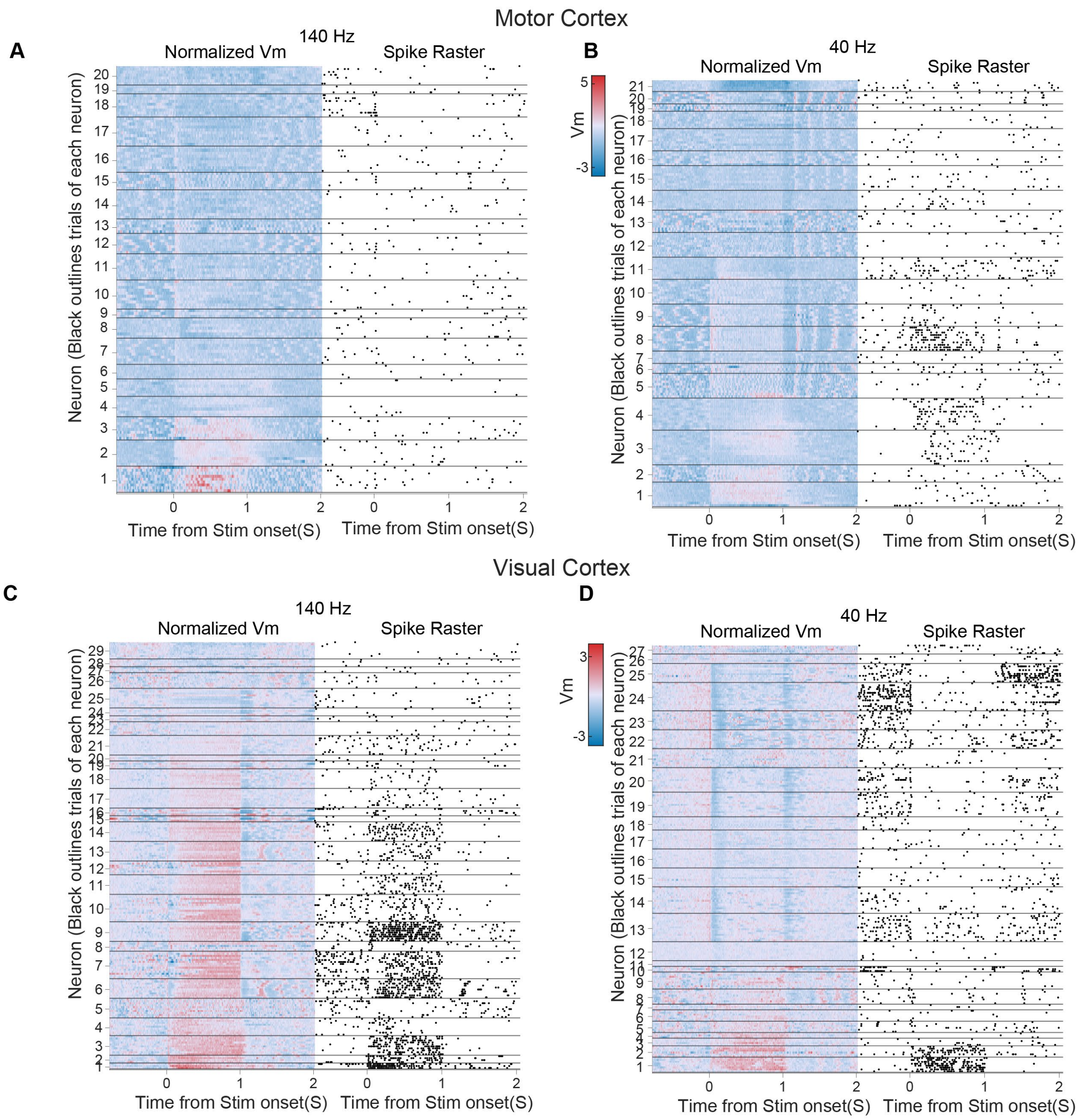


***Figure S3: Vm and spike raster of individual FSIs across stimulation trials****. (A) Vm heatmap (left) and corresponding spike raster (right) of FSIs in the motor cortex aligned to the onset of 140 Hz stimulation. Trials are separated for each neuron by horizontal black lines. (B) Same as a, but for 40 Hz stimulation. (C, D) Same as a, b, but for FSIs in the visual cortex.*

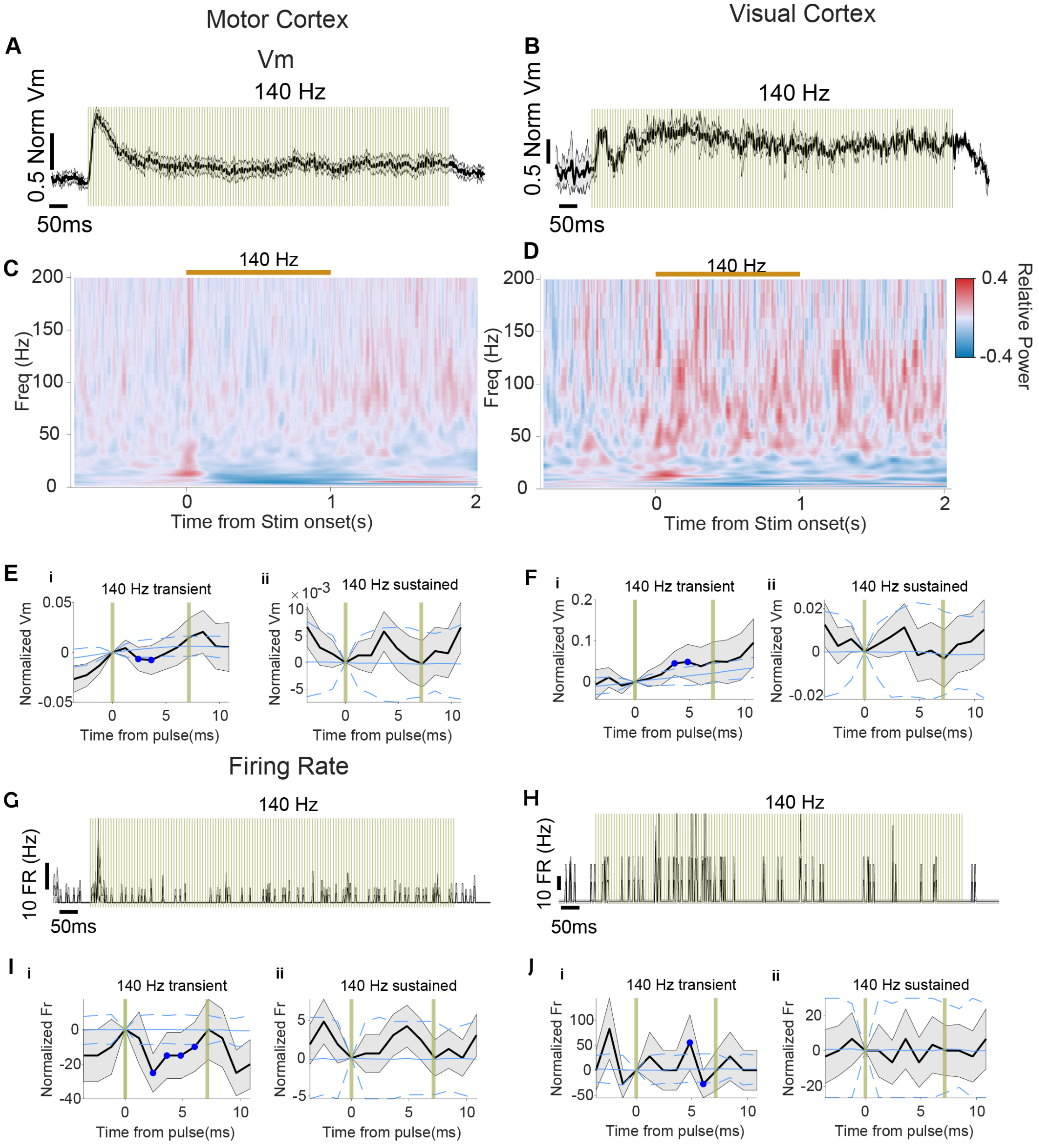


***Figure S4: Non-entrained FSIs as a population were weakly entrained by 140 Hz stimulation during the transient period of stimulation within 100 ms of onset.*** *(A) Population Vm across non-entrained motor cortex neurons (*n=11*) upon 140 Hz stimulation. (B) Same as a, but for FSIs in the visual cortex (*n=2*). (C) Vm power spectrum across non-entrained FSIs in the motor cortex upon 140 Hz stimulation. (D) Same as c, but for FSIs in the visual cortex. (E) (i) Population pulse-triggered Vm across non-entrained FSIs in the motor cortex during the transient period of 140 Hz stimulation. (ii) Same as e,i, but across pulses in the sustained period. Blue line indicates the shuffled distribution. Dashed lines indicate the 2.5% and 97.5% percentile of the shuffled distribution. Blue dots indicate points that are significantly different from the shuffled distribution. Gold vertical lines indicate individual pulses. Shaded gray area indicates SEM. (F) Same as e, but for FSIs in the visual cortex. (G) The population firing*

*rate across non-entrained FSIs upon 140 Hz stimulation. (H) Same as g, but for FSIs in the visual cortex. (I) (i) Population pulse-triggered firing rate of non-entrained FSIs in motor cortex during the transient period of 140 Hz stimulation. (ii) Same as I,i, but across pulses during the sustained period. (J) Same as I, but for FSIs in the visual cortex.*

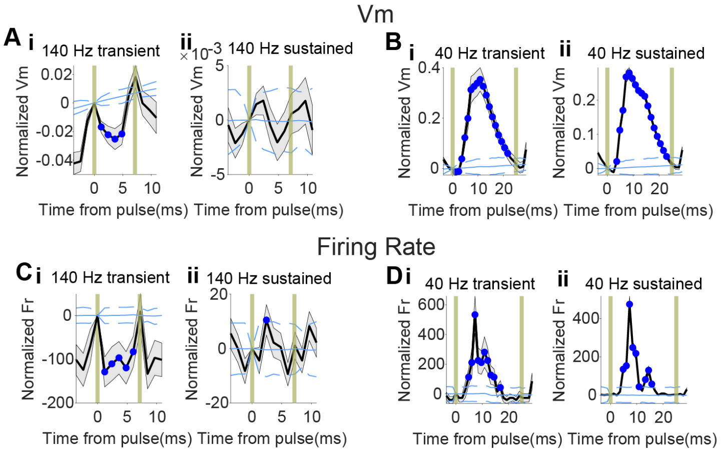


***Figure S5: CA1 pyramidal neurons' Vm was robustly entrained throughout 40 Hz stimulation, but only during the transient period of 140 Hz stimulation.*** *(A) (i) Population pulse-triggered Vm across CA1 pyramidal neurons during the transient period of 140 Hz stimulation (n=11). (ii) Same as a,i, but across pulses during the sustained period. (B) Same as a, but upon 40 Hz stimulation (n=13). (C) (i) Population pulse-triggered firing rate across CA1 pyramidal neurons during the transient period of 140 Hz stimulation. (ii) Same as c,i, but across pulses during the sustained period. (D) Same as c, but upon 40 Hz stimulation.*

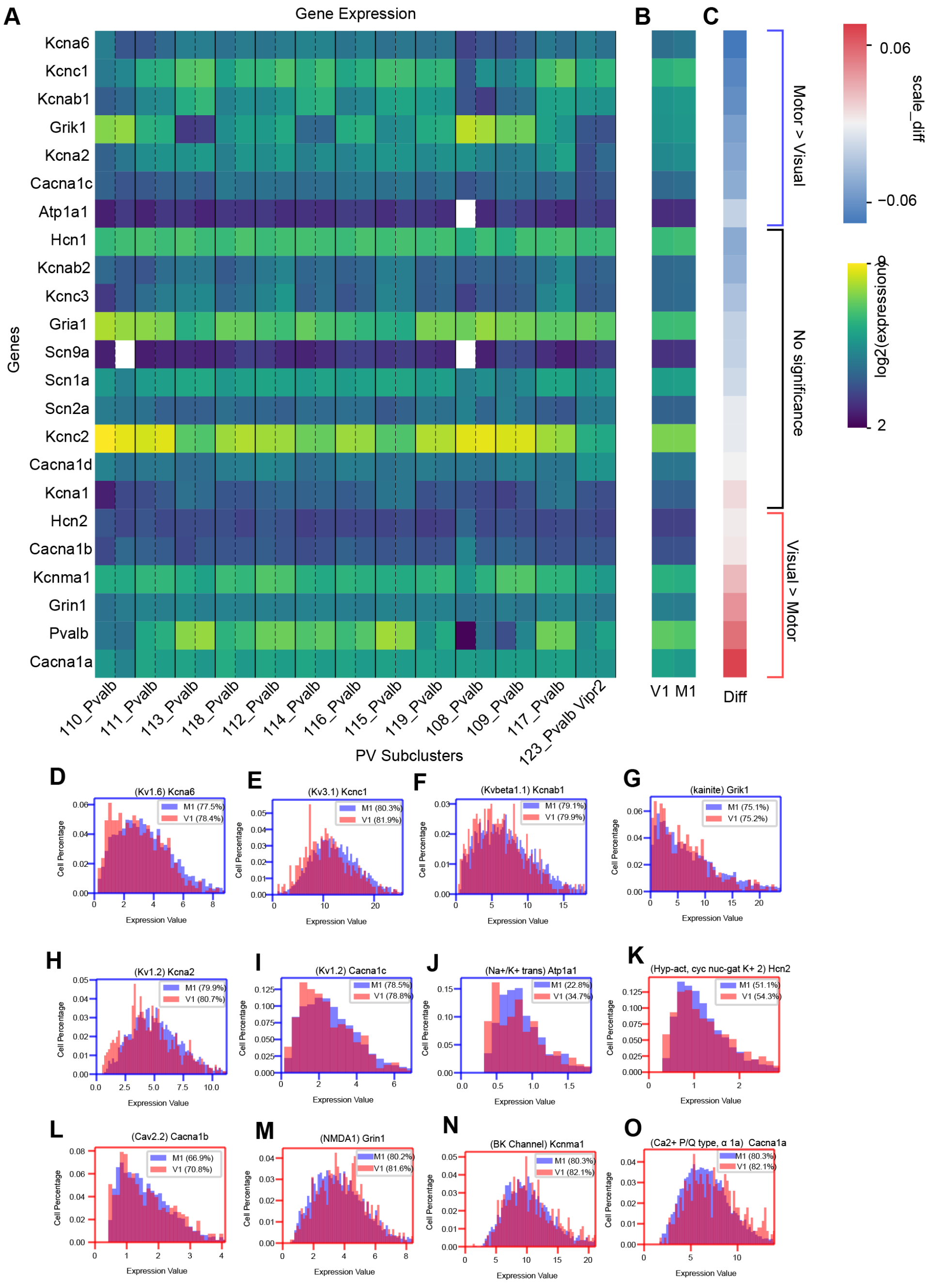


***Figure S6: Common ion channel expression levels for FSIs in the motor versus visual cortex.*** *(A) Heatmap of the relative expression values of common ion channels across subclusters of parvalbumin (PV) expressing FSIs (Shown are genes with significant expression determined using Wald test, p<0.05). Subclusters of PV cells in motor and visual cortices are separated by dotted black vertical lines. For each gene, the expression values were adjusted by the formula: (value – min(value))/(max(value) – min(value)). (B) Heatmap of mean expression value of common ion channels across all subclusters of FSIs in the visual and motor cortices (C)*

*Heatmap of the difference of ion channel expression between visual and motor cortex, calculated as ($value_{visual}$ – $value_{motor}$) / ($value_{visual}$ + $value_{motor}$). (D-J) Histogram of ion channels that expressed at a significantly higher level in the motor cortex than the visual cortex. (D) Kcna6 expression. Cliff's delta: 0.12; p=0.0096. (E) Kcnc1 expression. Cliff's delta: 0.18; p=0.0033. (F) Kcnab1 expression. Cliff's delta: 0.11; p=0.0028. (G) Grik1 expression. Cliff's delta: 0.074; p=0.013. (H) Kcna2 expression. Cliff's delta: 0.15; p=0.0004. (I) Cacna1c expression. Cliff's delta: 0.093; p=0.018. (J) Atp1a1 expression. Cliff's delta: 0.040; p=0.0068. (K) Hcn2 expression. Cliff's delta: 0.00061; p=0.048. (L-O) Histogram of ion channels that expressed at a significantly higher level in the visual cortex than the motor cortex. (L) Cacna1b expression. Cliff's delta: 0.013; p=0.011. (M) Grin1 expression. Cliff's delta: 0.076; p=0.042. (N) Kcnma1 expression. Cliff's delta: 0.066; p=0.012. (O) Cacna1a expression. Cliff's delta: 0.18; p=0.0.*

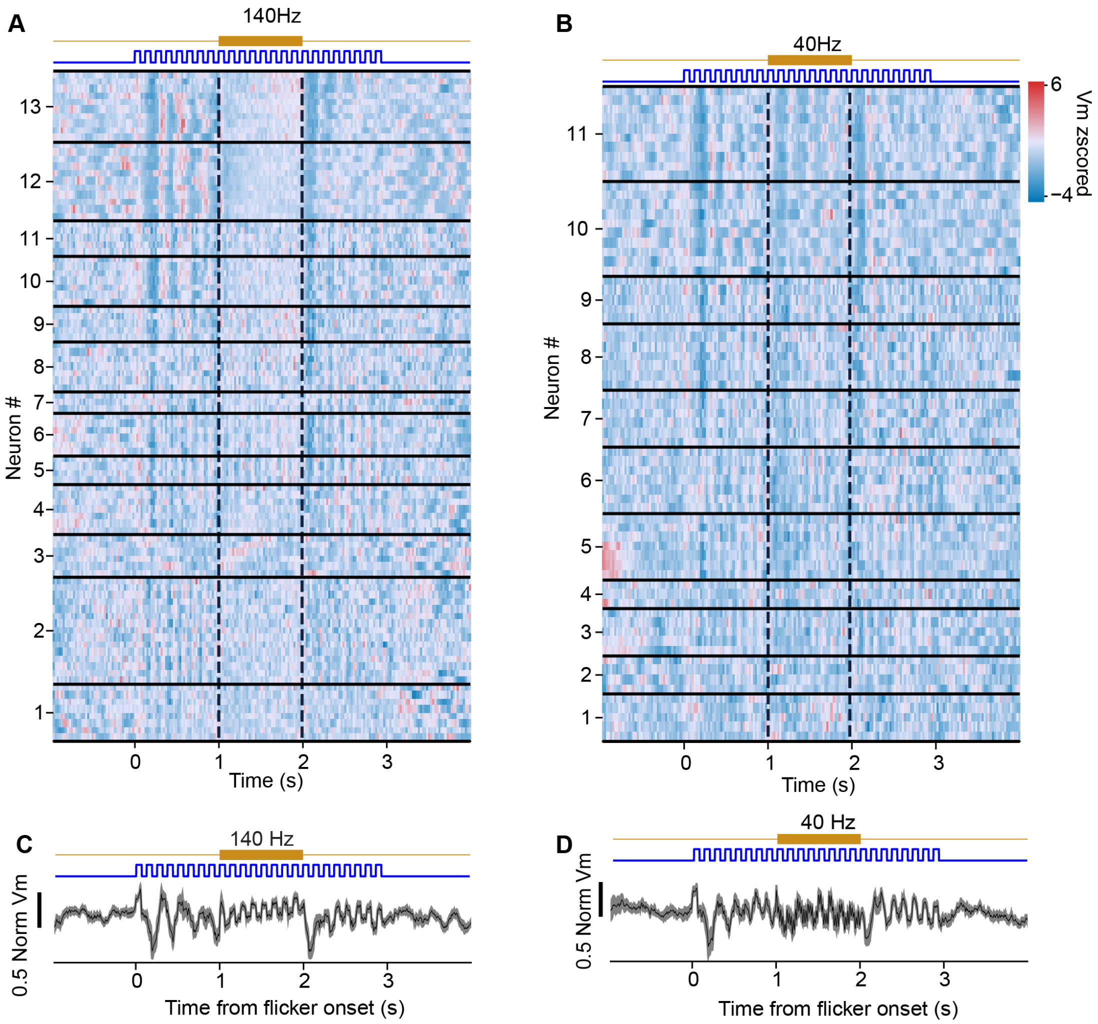


***Figure S7: Vm of individual visual cortex FSIs across stimulation trials during visual flicker presentation.*** *(A) Vm heatmap of FSIs upon 140 Hz stimulation during 8 Hz visual*

*flicker presentation. (B) Same as a, but upon 40 Hz stimulation. (C) Population Vm across FSIs upon 140 Hz stimulation. (D) Same as c, but upon 40 Hz stimulation. Shaded areas are the SEM.*

| | | Vm | | | Firing Rate | | |
|---|---|---|---|---|---|---|---|
| | | Transient | Sustained | Transient Vs Sustained | Transient | Sustained | Transient Vs Sustained |
| Visual Cortex | 140 Hz | 0.010622025 | 0.00049 | 0.0162709 | 0.01062 | 0.010622 | 0.761 |
| | 40 Hz | 0.229522943 | 0.22952 | 0.0422072 | 1 | 0.043285 | 0.1094 |
| Motor Cortex | 140 Hz | 0.004180908 | 0.21011 | 0.1961082 | 0.02127 | 0.210114 | 0.0023 |
| | 40 Hz | 0.000221252 | 0.0072 | 0.0495525 | 0.38331 | 0.078354 | 0.5901 |

***Extended Data Table 1**: This table lists the p-values from statistical tests for population Vm and firing rate changes. These tests yield significance for Figure 5.*

| | **140 Hz** | **40 Hz** |
|---|---|---|
| **Kruskal-Wallis** | 0.000437 | 0.5824 |
| **Dunn-Sidak** | | |
| Pre-stimulation vs stimulation | 1.74E-03 | 9.86E-01 |
| Pre-stimulation vs post-stimulation | 1.00E+00 | 8.60E-01 |
| Stimulation vs post-stimulation | 2.24E-03 | 6.72E-01 |

***Extended Data Table 2:** This table lists the p-values from statistical tests for power spectra during visual flicker period. These tests yield significance for Figure 6.*
